# A cross-validation-based approach for delimiting reliable home range estimates

**DOI:** 10.1101/168021

**Authors:** Eric R Dougherty, Colin J Carlson, Jason K Blackburn, Wayne M Getz

**Affiliations:** Department of Environmental Science, Policy, and Management, University of California, Berkeley, Berkeley, CA, USA.; Spatial Epidemiology and Ecology Research Laboratory, Department of Geography, University of Florida, Gainesville, FL, USA.; Emerging Pathogens Institute, University of Florida, Gainesville, FL, USA.; School of Mathematical Sciences, University of KwaZulu-Natal, Durban, South Africa.

**Keywords:** Time Local Convex Hulls, T-LoCoH, Home Range, Epidemiology, Visitation, Duration, Cross-validation

## Abstract

**Background:** With decreasing costs of GPS telemetry devices, data repositories of animal movement paths are increasing almost exponentially in size. A series of complex statistical tools have been developed in conjunction with this increase in data. Each of these methods offers certain improvements over previously proposed methods, but each has certain assumptions or shortcomings that make its general application difficult. In the case of the recently developed Time Local Convex Hull (T-LoCoH) method, the subjectivity in parameter selection serves as one of the primary impediments to its more widespread use. While there are certain advantages to the flexibility it offers for question-driven research, the lack of an objective approach for parameter selection may prevent some users from exploring the benefits of the method.

**Results:** Here we present a cross-validation-based approach for selecting parameter values to optimize the T-LoCoH algorithm. We demonstrate the utility of the approach using a case study from the Etosha National Park anthrax system. Utilizing the proposed algorithm, rather than the guidelines in the T-LoCoH documentation, results in significantly different values for derived site fidelity metrics.

**Conclusions:** Due to its basis in principles of cross-validation, the application of this method offers a more objective approach than the relatively subjective guidelines set forth in the T-LoCoH documentation and enables a more accurate basis for the comparison of home ranges among individuals and species, as well as among studies.

## 1 Background

Dramatic advancements in GPS telemetry devices have enabled researchers to gain a more comprehensive understanding of animal movement behaviors [1]. The decreasing costs of such devices have resulted in their widespread deployment and a capacity for data collection at unprecedented spatial and temporal resolutions [2]. Movement ecology has emerged as a discipline in its own right [3], with numerous methods and tools being developed and disseminated to analyze the wealth of available data. Ecologists can now quantitatively characterize home ranges and space use patterns over time. Often, the purpose of applying such quantification methods to movement paths is comparison of space use among individuals or species in order to examine such processes as niche partitioning [4, 5], optimal foraging [6, 7], social aggregation [8], or even decision-making [9]. However, many methods require user-defined input parameters, and results are often highly sensitive to the selection of such values. For meaningful comparisons, standardization is required [10], yet protocols to achieve consistency across applications are often non-existent.

One of the most fundamental concepts in movement ecology is the home range, conventionally defined as “the area traversed by the individual in its normal activities of food gathering, mating, and caring for young” [11]. Despite the apparent simplicity of this definition, the statistical formalization of the home range remains challenging, with alternative approaches emphasizing different aspects of animal movement and space use. The lack of a shared underlying theoretical framework makes comparison and standardization among methods all the more difficult, and the practical implications of selecting a particular conception of the home range make such considerations important.

Methods for home range delineation have evolved substantially since the concept of the home range first emerged in the literature [11]. The Minimum Convex Polygon (MCP) method was the most commonly used in the early years of home range description [12], despite its sensitivity to outliers [13] and its inability to further partition internal space [14]. The MCP-based conception of the home range lends itself naturally to some principles of space use in behavioral ecology, such as the general rule that individuals of territorial species often exhibit larger home ranges in relatively lower quality habitat. Kernel Density Estimation (KDE; [15]) emerged as a popular alternative that overcomes some of the limitations of the MCP method, but numerous parameter choices make comparisons among studies tenuous and replication of results dicult [16]. The KDE-based conception of the home range offers a probabilistic framing of animal space use, but may obscure some of the uncertainty inherent in movement data extracted at discrete time points. Both of these methods and their descendants also treat input points as independent, an assumption that is frequently violated with regularly sampled positions from movement paths. E orts to overcome this inherent autocorrelation have included resampling or weighting algorithms [17, 18], but more recently, methods like the Brownian Bridge Movement Model (BBMM; [19]) and autocorrelated KDE (AKDE; [20]) have been developed to explicitly incorporate the serial nature of movement data. These more nuanced conceptions of the home range and movement behaviors account statistically for uncertainty and autocorrelation, but reliance on random walk dynamics and related assumptions may not account for the behavioral dependency of animal movements [21]. While some of the earlier home range delineation methods could be built for multiple individuals simultaneously, many of these more rigorous methods are parameterized for each individual separately.

The recently developed Time Local Convex Hull method (T-LoCoH; [22]) builds upon the non-parametric LoCoH method [23] by explicitly integrating the temporal component of movement data, effectively scaling time with distance in the construction of local point sets, or hulls. Essentially, this method is governed by a simpler, MCP-based conception of the home range, but works at a finer spatiotemporal scale and enables extension to a more probabilistic description of space use. The T-LoCoH algorithm constructs a utilization distribution (UD) by aggregating local convex polygons, or hulls, built around each point. The hulls are created by selecting the *k* nearest neighbors of a given point and then sorted by density and merged together to form the UD. The selection of nearest neighbors can be modified by the inclusion of a dimensionless scaling parameter *s*, which transforms the time interval between points into a third axis in Euclidean space. The distance between points in this three-dimensional volume is called time-scaled distance (TSD), and it serves to separate points that are far apart in time despite their close proximity in two-dimensional space. Thus, an *s* value of zero will produce the same home range as the original LoCoH method. Guidelines exist for choosing appropriate values to construct a suitable home range, but much discretion is left to the researcher based on the particular subject of their inquiry [22].

A similar approach relies upon the parameter *a*, which selects nearest neighbors whose distance from the focal point sums to the value *a*. This method also requires the *s* parameter for weighting the TSD, but the alternative parameterization may be especially useful for more adaptive hull creation, such that more densely clustered areas of the movement path result in hulls with more points than areas of sparse usage [22]. A rough sensitivity analysis reveals that small differences in either of these parameters has dramatic impacts on the qualities of the resulting home range. The values of these parameters are also contingent upon the movement path itself, meaning that the paths of individuals of the same (or different) species may not result in comparable home ranges. To make such comparisons ecologically and statistically sound, the procedure must be standardized, but to date no such method exists.

Here we demonstrate the use of a novel cross-validation-based method to optimize parameter value selection for implementing the T-LoCoH algorithm based on the unique qualities of each individual movement path. This approach overcomes much of the subjectivity inherent in the recommended parameter selection protocol [22], circumventing the primary challenge to building and interpreting T-LoCoH home ranges. In addition, this method has the added benefit of enabling comparisons of home range features and derived metrics across individuals, species, and spatiotemporal scales, as the same underlying characteristics are used to select the optimal parameter values. We demonstrate the utility of this method with a case study on herbivore movement in the anthrax-dominated landscape of Namibia’s Etosha National Park.

## 2 Methods

### 2.1 Case Study

Pathogens indirectly transmitted via environmental reservoirs (e.g., water, soil, or animal excretions) represent a unique challenge for ecologists and epidemiologists. Risk of infection in such cases will depend upon the particular conditions at reservoirs [24, 25], the feeding behavior of the host [26, 27, 28], and the spatial arrangement of reservoir sites relative to susceptible animals [29], all of which may serve to facilitate or dilute pathogen transmission. Certain characteristics of movement behavior may aid in identifying the variation in risk of infection among individuals of the same and different species, including home range size [30], site fidelity [31, 32], and contact network structure [33, 34]. Comparisons of movement-associated transmission risk across individuals may serve to guide management e orts in areas affected by environmentally borne pathogens by identifying high-risk individuals and areas [35, 36], but a failure to explicitly account for individual differences may preclude robust evaluations of epidemiologically-relevant space use patterns [37]. We applied our novel method to the movement trajectories of individuals from two herbivore species in relation to anthrax (the acute disease caused by the bacterium *Bacillus anthracis*) in Etosha National Park, Namibia. As a disease transmitted via environmental reservoirs, anthrax represents an ideal case study for exploring the connections between individual movement on the landscape and resulting disease risk.

GPS point locations were obtained for individuals of two different susceptible ungulate species during the anthrax season in Etosha National Park, Namibia. For both the plains zebra (*Equus quagga*) and springbok (*Antidorcas marsupialis*), the anthrax season was defined as the five-month period between February 1 and June 31 [36]. Due to differences in the temporal resolutions at which the data were initially collected, subsets of the data were created so that each individual had one point location per hour throughout the sampling period. The total number of points for each individual during this period ranged from 2111 to 3601 (Table 1). Any missing data values during the sampling period were estimated using a Kalman smoothing approach [38]. Plains zebra and springbok show no sex-related disparity in infection rate [39]. All five zebra individuals chosen for analysis were female, while four of the six springbok were female and two were male.

**Table 1.**
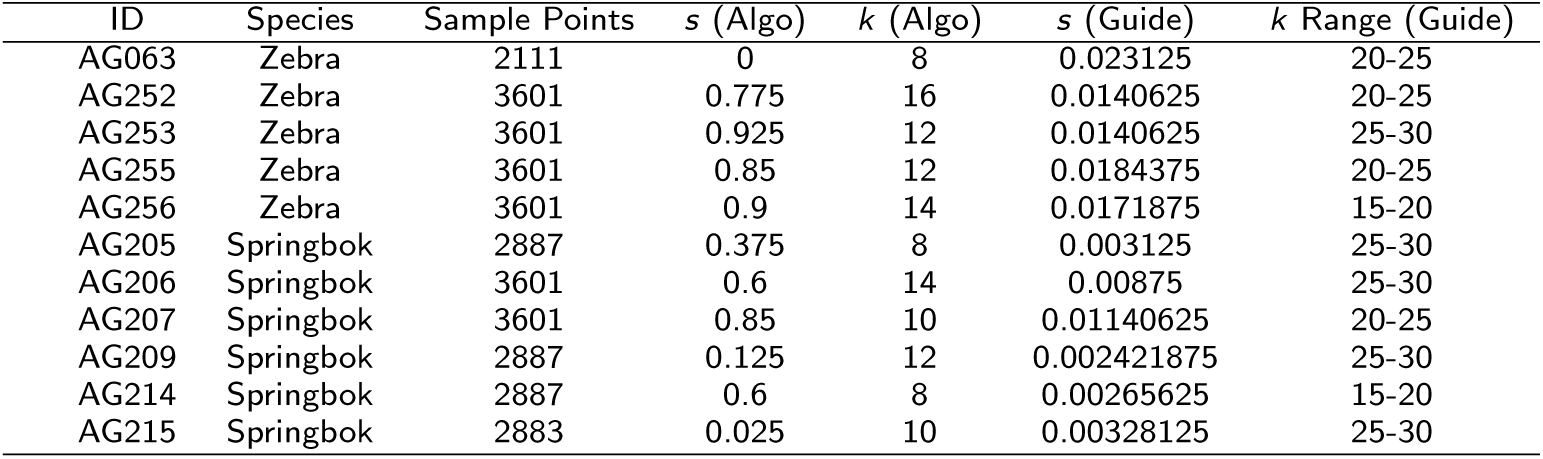
Parameter values for analysis. The *s* and *k* values selected using the algorithm and the guidelines in the T-LoCoH documentation. A range of *k* values were used for the Guide due to the subjective nature of parameter selection.

### 2.2 Existing Parameter Selection Protocol

The *k* (number of nearest neighbors) and *s* (time-scaled to distance) parameter values obtained using the proposed algorithm (below) were compared to those one might select based on the guidelines set forth in the T-LoCoH documentation [22]. In addition, the derived metrics, including visitation rate (the number of visits to a given hull, separated by a pre-defined amount of time) and mean duration (the average number of relocations within a hull during each of those visits) were compared to determine the impact of selecting these alternative parameter sets on epidemiologically and ecologically meaningful measures. Because these values are calculated at the scale of the hull, they are likely to strongly depend upon the size of the hulls themselves, with larger hulls leading to relatively higher duration and lower visitation rates as it becomes more difficult to “leave” a hull. The selection of values for the *k* and *s* parameters will therefore have implications on the mean values calculated for each individual.

To select appropriate *k* and *s* values using the guidelines, the proportion of time-selected hulls (PTSH) method was used. The PTSH approach calculates the distances between pairs of points under a set of alternative *s* values, and notes the proportion of pairs that are selected due to their temporal proximity rather than their spatial proximity. Ten repetitions of the method were implemented for each trajectory and all *s* values associated with a PTSH between 0.4 and 0.8 were obtained from each run. The median value was then chosen from this set and assigned as the *s* value for that individual. Using these *s* values, six potential isopleth sets were created, ranging from *k*=5 to *k*=30 in increments of 5. Isopleths are created after the hulls are merged together by taking their union, whereby the *i^th^* isopleth contains i-percent of points. The *k* values used in subsequent analyses were chosen using two independent researchers who were asked to select an isopleth set (or range of sets) that satisfied the minimum spurious hole covering criteria, which calls for the selection of the smallest *k* value that minimizes the holes present in the core area of the individual’s home range. To convey the subjectivity associated with the *k* selection procedure, both the lower and upper bounds of the ranges of *k* values selected by the independent researchers were mapped and derived metrics extracted.

### 2.3 Cross-Validation-based Parameter Selection

In developing a cross-validation-based approach to parameter selection, we aim to remove much of the subjectivity in the process and enable the data to inform appropriate values. The cross-validation method depends upon the creation of a series of training and testing data sets. For each set, test points were chosen randomly from the full movement path such that approximately one out of every 450 sampled points was selected as a test point (thus, each point had a probability of 0.002222 of being a test point). To ensure independence of the testing points, the 50 points preceding and following each selected test point were removed from the full dataset, and the remainder was considered the training data. For a path with 3600 points, this results in the selection of 8 test points, on average, for each testing set, leaving 2792 points in the training set. The resulting training datasets therefore consisted of approximately 80% of the original data points (Figure 1). To minimize variation in the procedure, this stochastic splitting process is repeated *n* times (in this case, 100) for each movement path.

**Figure 1.**
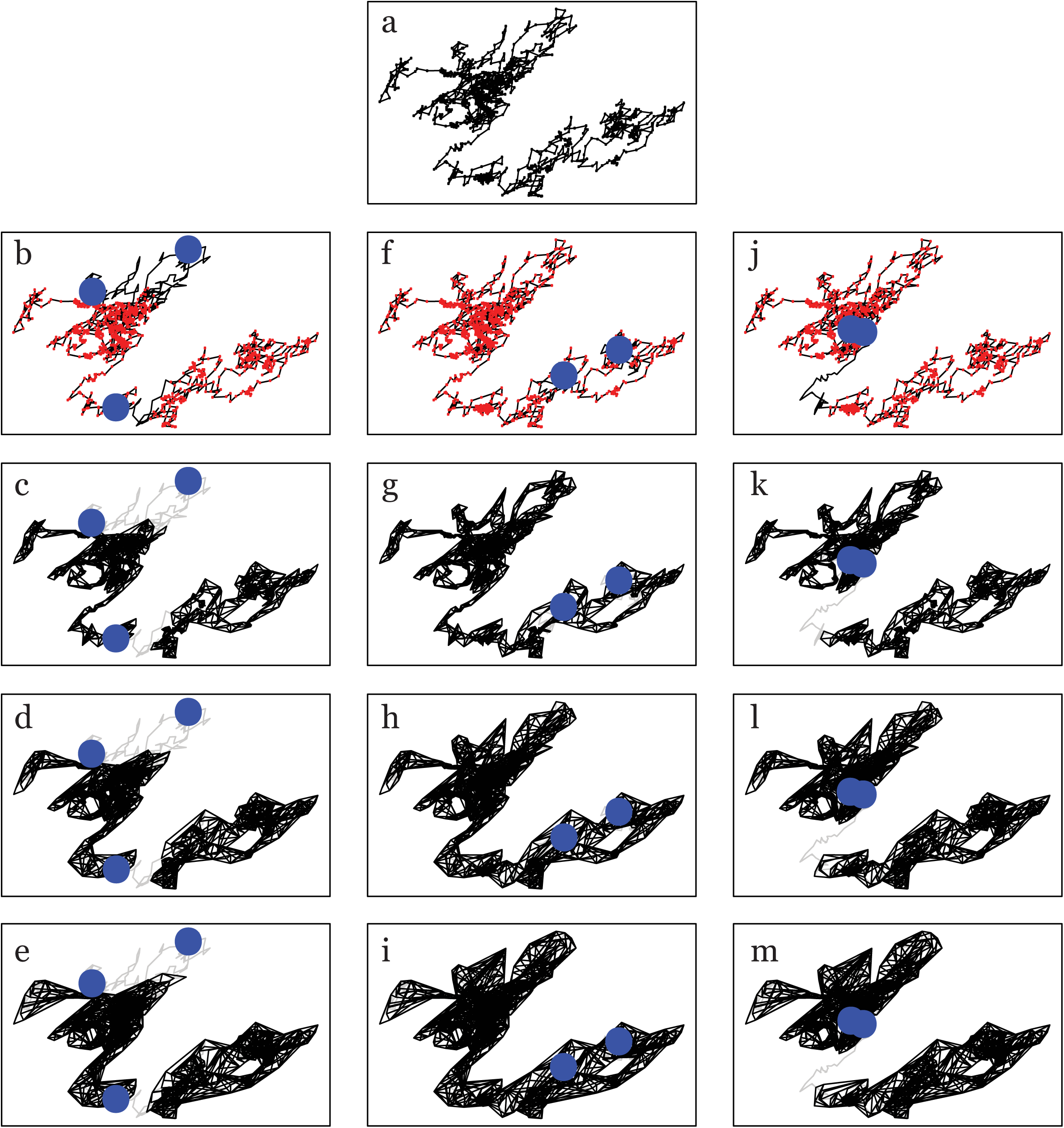
Conceptual Figure of the Proposed Algorithm. A test case of the algorithm using a simulated movement trajectory of 1000 relocation points (a). Three of the subsets of those points, with red points indicating those locations that remain in the training sets and blue points representing the test points for the later probability calculation (b,f,j). For each subset of points, a hullset is created using T-LoCoH, with an arbitrarily chosen *s* value of 0.5 and *k* values of 5 (c,g,k), 15 (d,h,l), or 25 (e,i,m). These three subsets serve to illustrate three possible scenarios as the *k* values increases: either test points that are not covered by the hull set at low *k* values continue to be uncovered with high *k* values (left-hand column), test points that were not originally covered by the hull set at smaller *k* values becomes covered (center column), or test points are covered at low *k* values and continue being covered at higher *k* values (right-hand column).

A grid-based exploration of parameter space was then conducted (Figure 2), whereby each of the 100 training/testing datasets was analyzed at every combination of *k* and *s* values on the grid. This analysis entailed the creation of local convex hulls with *k* nearest neighbors and a scaling factor of *s*. In all subsequent analyses, we assume that the scaling of time follows a linear formulation; however, when movement patterns more closely exemplify di usion dynamics, an alternative equation for the TSD may be more accurate [22]. The test points were then laid upon the resulting hulls, and the probability of each was calculated as the proportion of the total number of hulls (equivalent to the total number of points in the training dataset) that contained the test point (Figure 1). Test points that were not contained within any hulls were assigned a probability equal to the inverse of the total number of points in the full movement path divided by 100, effectively penalizing any hull sets that did not include each of the test points. Though an arbitrary selection, the choice of a consistent penalty term across individuals will serve to standardize the procedure. A larger penalty will likely result in a higher optimal *k* value and bear a closer resemblance to the MCP. The natural log of the probability was calculated and information criterion values analogous to Akaike’s Infromation Criterion (AIC) were derived using the equation:

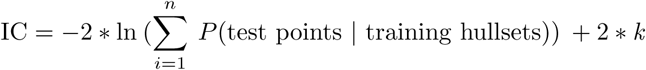

**Figure 2.**
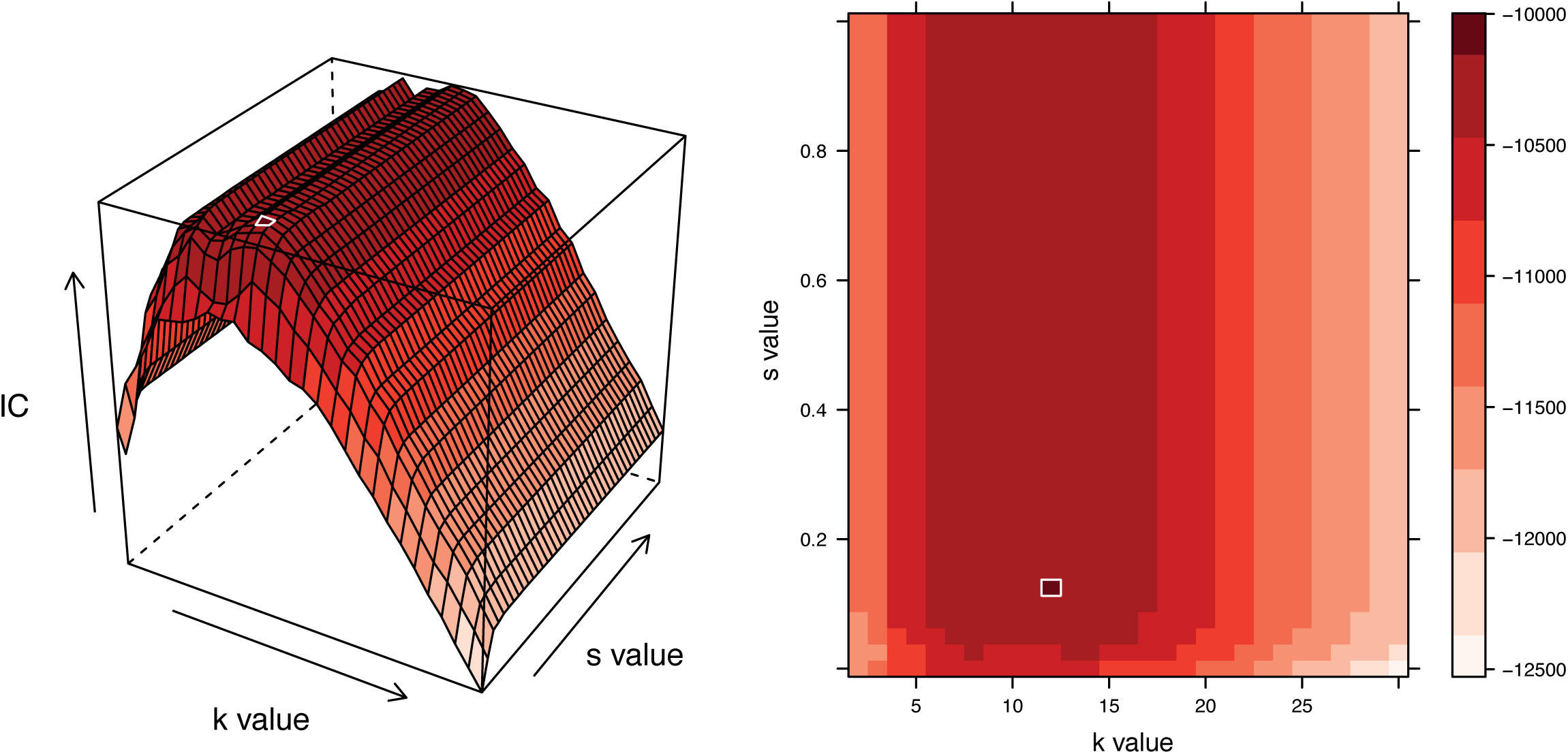
Conceptual Figure of Grid-based Search. An information surface is generated as the algorithm searches over a grid of alternative *s* and *k* values for each individual movement path. The increments of the grid can be chosen by the user. The peak in the surface indicates that the maximum level of information about the home range is obtained at the associated parameter set. The ridge in surface illustrated here suggests that multiple parameters sets with different *s* values offer the same information content. In such cases, the parameter set with the smallest *s* and *k* values was selected. Here, the white outlines denote the highest information criterion value, and thereby, the optimal parameter set.

The choice of 2*k* as the penalty term was made to maintain a structure analogous to the AIC equation. Given the expansive literature concerning the performance and behavior of AIC under various scenarios, maintaining this structure may insight into similar strengths and weaknesses of the proposed approach. Ultimately, without such a penalty, all movement paths would tend towards a *k* equal to the number of points in the training set, such that each individual point was assigned a probability of one. It should be noted that this penalty term is specific to the *k* (nearest neighbors) method, but the underlying cross-validation procedure could very easily be extended for the optimization of the *a* (adaptive parameter) method if an appropriate penalty term is selected. An ideal penalty term would likely result in a increase of the information criterion value by a similar magnitude as in the *k*-based formulation above (i.e., ranging from approximately 10^0^ to 10^2^).

Despite the use of a testing and training dataset in the creation of the hullsets, we deemed that the use of a measure of sensitivity versus specificity, such as the receiver operator characteristic (ROC) curve, would not serve as an effective means of comparing alternative parameter sets. While false negatives (i.e., test points that are not contained within any hulls) are certainly easy to measure, without some form of pseudo-absence point, one cannot easily obtain a false positive rate (i.e., points that fall within the home range defined by the hulls, but not actually a point occupied by the animal). Rather, the log probability measure was chosen, as test points can be penalized for being false negatives by assigning a consistent small value as its probability, but there is no need to create pseudo-absence points or account for false positives in any way.

The grid-based search of parameter space allows for the identification of the combination of *s* and *k* values that offer the optimal information criterion value (Figure 2). In the case that multiple *k* or *s* values offer the same level of information, a ridge will appear in the information content surface. This was not especially uncommon in the paths analyzed here, particularly along the *s* value axis. This likely indicates that relatively large differences between the optimal *s* value from the proposed algorithm and that derived using the guideline-based criteria may reflect relatively small differences in actual information content. Further, this common shape to the resulting surfaces suggests that the *k* value tends to have a much more dramatic impact on the criterion. R code for a parallelized version of the algorithm using this grid (where each individual movement path is run in parallel) is supplied in the supplementary materials.

## 3 Results

The algorithmic grid-based search covered *s* values from 0 to 1 in increments of 0.025 and *k* values between 2 and 30 in increments of 1. In the subsequent statistical analyses, the results of paired *t*-tests are presented to demonstrate the significance of differences when the proposed method was used relative to the guide-based parameter selection criteria, beginning with the *k* and *s* parameters themselves (Table 1). The mean *k* value selected using the algorithm for springbok (*N* = 6) was 10.33 (SE = 0.95) and for zebra (*N* = 5) was 12.40 (SE = 1.33). The median of the range of *k* values selected using the T-LoCoH guidelines was used for comparison with *k* values resulting from the use of the algorithm. The mean of these median values was 22.5 (SE = 1.71; *p* < 0.001) for the springbok and 20 (SE = 1.58; *p* = 0.03) for the zebra. The mean *s* value selected using the algorithm for springbok was 0.43 (SE = 0.13) and for zebra was 0.69 (SE = 0.17). The mean *s* value selected using the guidelines was 0.0053 (SE = 0.0016; p = 0.02) for springbok and 0.017 (SE = 0.0017; *p* = 0.02) for zebra. These results indicate that significantly different *s* and *k* values are obtained when using the algorithm rather than the upper or lower *k* and *s* values from the guidelines. According to either method, the difference in the *s* values between species indicates that zebra move greater distances than springbok in the same amount of time, further supported by other qualities of the home range.

In terms of the area of the home ranges resulting from each parameter set (Table 2), comparisons were conducted using both the low and high values from the range of the guideline-based parameters relative to the algorithm-based parameter set. The mean home range area for springbok using the algorithm was 203.88 km^2^ (SE = 59.71 km^2^) and 602.64 km^2^ (SE = 116.77 km^2^) for zebra. The mean home range area for springbok using the low value of the range based on the guidelines was 251.22 km^2^ (SE = 72.51 km^2^; *p* = 0.04) and 660.84 km^2^ (SE = 74.30 km^2^; *p* = 0.42) for zebra. The mean home range area for springbok using the high value of the guideline-based range was 265.41 km^2^ (SE = 76.23 km^2^; *p* = 0.03) and 694.43 km^2^ (SE = 80.81 km^2^; *p* = 0.21) for zebra. This indicates that the home ranges constructed using the algorithm-based parameter set were significantly lower than even the low value in the guideline-based range for springbok, but the difference between the resulting home ranges using either set from the guidelines and the algorithm were not statistically significant for the zebra.

**Table 2.**
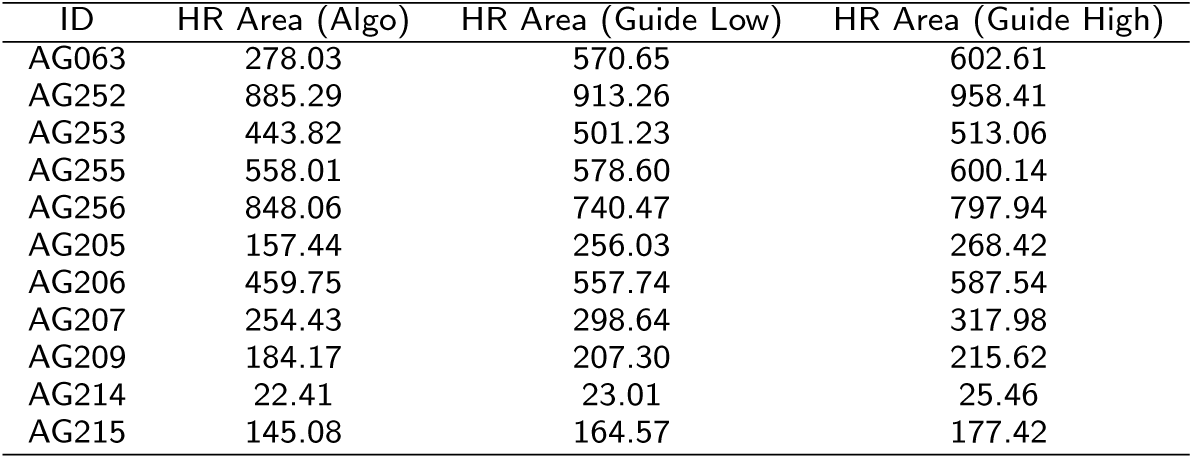
Home range areas (in square kilometers). The total area of the home range obtained using the parameter sets recommended by the algorithm and by the guidelines set forth in the T-LoCoH documentation.

For the derived fidelity metrics, duration (Table 3) and visitation (Table 4), comparisons only concern the mean values of each metric for each individual, though other descriptive statistics of the distribution of all duration and visitation values may be of interest in some cases. The mean duration (MNLV) for springbok using the algorithm values was 13.39 (SE = 1.84) and for zebra was 7.52 (SE = 0.96). Mean duration derived using the low values in the range of *s* and *k* values obtained based on the guidelines were between 21.47 (SE = 3.84; *p* = 0.01) for springbok and 9.72 (SE = 0.47; *p* = 0.11) for zebra. The mean duration derived using the high values in the guideline-based range were 24.35 (SE = 4.20; *p* = 0.007) for springbok and 11.11 (SE = 0.49; *p* = 0.03) for zebra. The mean visitation rate (NSV) for springbok using the algorithm values was 7.37 (SE = 1.92) and 7.58 (SE = 1.65) for zebra. Mean visitation rates derived using the low value from the range of *s* and *k* values obtained using the guidelines were 8.38 (SE = 2.06; *p* = 0.01) for springbok and between 8.39 (SE = 1.71; *p* = 0.36) for zebra. Using the high value from the guideline-based range, the mean visitation rate is 9.00 (SE = 2.27; *p* = 0.008) for springbok and 9.40 (SE = 1.77; *p* = 0.07) for zebra. Once again, the derived metrics exhibit the patterns observed in the home range area comparison; the use of the algorithm results in statistically significant changes in the mean duration and visitation rates for the springbok, but not for the zebra.

**Table 3.** Mean duration (MNLV) values. The derived metrics obtained using the parameter sets recommended by the algorithm and by the guidelines set forth in the T-LoCoH documentation.

| ID | MNLV (Algo) | MNLV (Guide Low) | MNLV (Guide High) |
| --- | --- | --- | --- |
| AG063 | 4.07 | 10.02 | 11.32 |
| AG252 | 9.88 | 10.37 | 11.74 |
| AG253 | 7.64 | 10.71 | 12.45 |
| AG255 | 7.57 | 9.50 | 10.34 |
| AG256 | 8.42 | 8.00 | 9.70 |
| AG205 | 13.24 | 24.38 | 27.10 |
| AG206 | 10.92 | 14.32 | 16.42 |
| AG207 | 7.95 | 12.26 | 14.47 |
| AG209 | 16.74 | 23.41 | 26.04 |
| AG214 | 11.10 | 16.54 | 19.43 |
| AG215 | 20.39 | 37.89 | 42.63 |

**Table 4.** Mean visitation (NSV) values. The derived metrics obtained using the parameter sets recommended by the algorithm and by the guidelines set forth in the T-LoCoH documentation.

| ID | NSV (Algo) | NSV (Guide Low) | NSV(Guide High) |
| --- | --- | --- | --- |
| AG063 | 2.74 | 5.82 | 6.58 |
| AG252 | 6.01 | 5.64 | 6.30 |
| AG253 | 12.82 | 15.00 | 16.04 |
| AG255 | 7.84 | 8.07 | 9.46 |
| AG256 | 8.52 | 7.43 | 8.62 |
| AG205 | 2.77 | 4.24 | 4.50 |
| AG206 | 6.07 | 6.46 | 6.90 |
| AG207 | 12.92 | 14.85 | 15.67 |
| AG209 | 3.28 | 3.60 | 3.80 |
| AG214 | 13.52 | 14.56 | 16.3 |
| AG215 | 5.64 | 6.57 | 6.83 |

## 4 Discussion

The concept of the home range remains a contentious one, with some researchers suggesting that the choice of delineation method should be defined by the question at hand [10]. When comparison is an element of an analysis, however, standardization of sampling protocols and estimation techniques is required [37, 14]. Considering the multitude of statistical issues overcome by the T-LoCoH method, it should become an increasingly prevalent tool for such analyses. Therefore, eliminating subjectivity from the procedure represents an important step for enabling comparisons both within and among species and studies.

One important consideration is that the “true” *k* and *s* values are inherently unknowable. Even the use of simulation methods, which would o er perfect knowledge of the position of an agent at any given time, would not enable the construction of a “true” home range because that would entail the selection of one particular conception of the home range. The approach laid out here offers one such conception, where consistency, as measured by the ability to capture testing points in home ranges created using a subsample of the full movement trajectory, is valued above other measures, such as contiguity or inclusion. By applying this conception of the home range to movement data from different individuals or species, the proposed method effectively unifies the resulting home ranges, enabling further comparison.

Recent empirical studies utilizing the T-LoCoH algorithm for delineating home ranges illustrate the subjectivity involved in parameter value selection [40, 41, 42]. While many studies rely upon the guidelines set forth in the Tutorial and Users Manual provided by the creators of the ‘tlocoh’ package in R [22], there was some variation among studies regarding the selection of *s* values (i.e., choosing different proportions of hulls that are considered time-selected) and whether the *k* or *a* approach was used for selecting nearest neighbors. Most of the home range studies applying the T-LoCoH method do so across multiple individuals, and researchers must decide whether to select separate parameter values for each individual or to have a single overarching parameter set. This decision is particularly important in cases where multiple species are being compared [43], as attribution of differences in home ranges to actual ecology rather than parameter choice may be muddled. Most troubling, however, is the fact that several studies implementing T-LoCoH neglect to specify the parameter values they ultimately used for their analyses, making replication of results nearly impossible.

With regard to the decision about a single parameter set used across individuals or separate sets for each movement path, we argue that consistency and comparability does not emerge from the parameter sets themselves. Rather, the resulting home ranges can be unified by the home range conception that guided their creation. As previously mentioned, the method proposed here serves as that unifying conception, prioritizing consistency in the home range through the use of a cross-validation approach. In order to construct such a home range for a particular individual, a very different parameter set from another individual may be necessary. Thus, we recommend the use of the proposed algorithm (and the underlying conception of the home range upon which it is built) to make home range analyses more readily comparable between movement tracks.

The results from this case study indicate several important trends. The first is that the *s* and *k* parameter sets selected by the cross-validation-based approach are significantly different from those one would obtain using the proportion time-selected hulls (PTSH) method and the minimization of holes approach set forth in the T-LoCoH documentation. While the guidelines may be suitable in some cases, the average difference between the algorithm-based *k* value and the lower bound of that chosen using the guidelines is over 10, suggesting that using isopleths to judge a home range rather than the hulls results in higher *k* values. Similarly, the *s* values selected by the algorithm are orders of magnitude larger than those selected by the PTSH method, demonstrating the important role of the temporal aspect in predicting space use patterns in the proposed approach.

Another important trend concerns the relationship between the derived metrics of visitation and duration. While both are ecologically and epidemiologically important measures of individual space use patterns that rely upon the underlying hullsets, the mean visitation value was not statistically significantly different when the algorithm was used rather than the guidelines while the mean duration did exhibit marked differences. This suggests that the two metrics respond differently to changes in *k* and *s* values, a somewhat surprising result given that one might expect a simultaneous increase in both values, or a trade-off, such that an increase in one would indicate a decrease in the other. This non-parametric scaling further suggests the importance of having a standardized method for selecting *k* and *s* values.

Changes in site fidelity metrics can have important ecological implications. For diseases like anthrax, which are caused by indirect pathogen transmission at environmental reservoirs, if a locally infectious zone (LIZ; [44]) is present within the home range of an individual, a greater level of site fidelity is likely to place the individual at repeated and extended risk of encountering the pathogen. However, this same high site fidelity may protect an individual against exposure if there are no LIZs in the home range. Consequently, higher mean visitation and duration values are likely to produce a greater level of heterogeneity of infection risk for individuals within a spatially structured population.

In the case of these particular herbivores in the Etosha system, this difference in heterogeneity may be observed in the relative likelihood of a lethal versus non-lethal infection in the two species. The zebra population in Etosha is approximately 13,000 and the springbok population is estimated at 15,600 [44]. After accounting for imperfect detection [45], carcass surveillance data from 2000-2013 suggest that the mean annual mortality rate directly linked to anthrax is approximately 1.34% (95% CI: 0.80% - 1.88%) in zebra and 0.26% (95% CI: 0.18% - 0.35%) in springbok. Additionally, the rate of sub-lethal exposure as indicated by the existence of antibodies in blood serum samples is between 52% and 87% for zebra and between 0% and 15% for springbok [44]. Based on the high values of non-lethal infection, the annual rate of a zebra exposed to anthrax experiencing a lethal dose is approximately 1.5% whereas exposed springbok experience a lethal dose at an annual rate of approximately 1.8%. This suggests that the zebra population may experience higher overall exposure rates to the pathogen, but because of their relatively low mean duration, a large proportion of the exposed population will contract a non-lethal dose, as they will move on from LIZs relatively quickly. The greater mean duration value observed in the springbok population would lead to expectations that some individuals will experience high doses based on repeated and lengthy visits to LIZs or no exposure, with moderate, non-lethal exposure being fairly rare.

The same principles can be applied to other disease systems, where indirect pathogen transmission may be linked to the spatial overlap of a species shedding a pathogen into the environment and naive hosts of another species contacting the pathogen during commingling, as in the case of brucellosis [46]. Commingling, frequently calculated as a function of home range overlap, is a common measure of inter-specific transmission risk, particularly between livestock and wildlife (e.g., bovine tuberculosis [47, 48]). The use of the algorithm enables the construction of comparable home ranges among different species with greater confidence, thereby overcoming one of the most important challenges of using and interpreting T-LoCoH and allowing for a broader application in multi-species disease systems. Though the general patterns observed using the guideline-based parameter sets are similar to those observed using the algorithm-based parameter sets, there are some potentially important differences regarding a derived metrics of space usage. Namely, the difference in the mean duration values between the two species has decreased in magnitude. While the difference in the mean duration values of the species remains significant, the algorithm-based parameter set results in values that suggest a greater level of similarity between the two species than those obtained using the guidelines to select the *s* and *k* values. These subtle differences may ultimately have significant consequences for the overall accuracy of agent-based models in ecoepidemiology, and a more consistent method of selection can ultimately accelerate the development of those models for such systems.

Finally, the concept of the probabilistic home range was an important advancement in the home range literature [16], but in the case of T-LoCoH, where isopleths are built atop a series of hulls, the resulting home range may represent an over-fitting to the data (Figure 3b). As such, this process may be useful for identifying core areas, but may overlook corridors or treat such outlying landscape features as part of the core area by altering the parameter set to fill in “holes” in the home range. The guidelines aim to minimize holes in the core area of the home range, but because they are based on the probabilistic isopleths, the hulls may need to grow considerably (i.e., the *k* value must increase) before the underlying hulls predict presence in those areas. Using the hulls underlying those isopleths themselves may represent an underfitting to the data (Figure 3e), in essence, a return to the MCP concept whereby too much unused space would be considered suitable. The algorithm circumvents the intermediate step of using isopleths by minimizing holes in the hullset itself. The home range that one builds from these hulls (based on a considerably smaller *k* value) may therefore represent an ideal trade-off between the overfitting of the isopleths and the underfitting of the hulls at an inflated *k* value.

**Figure 3.**
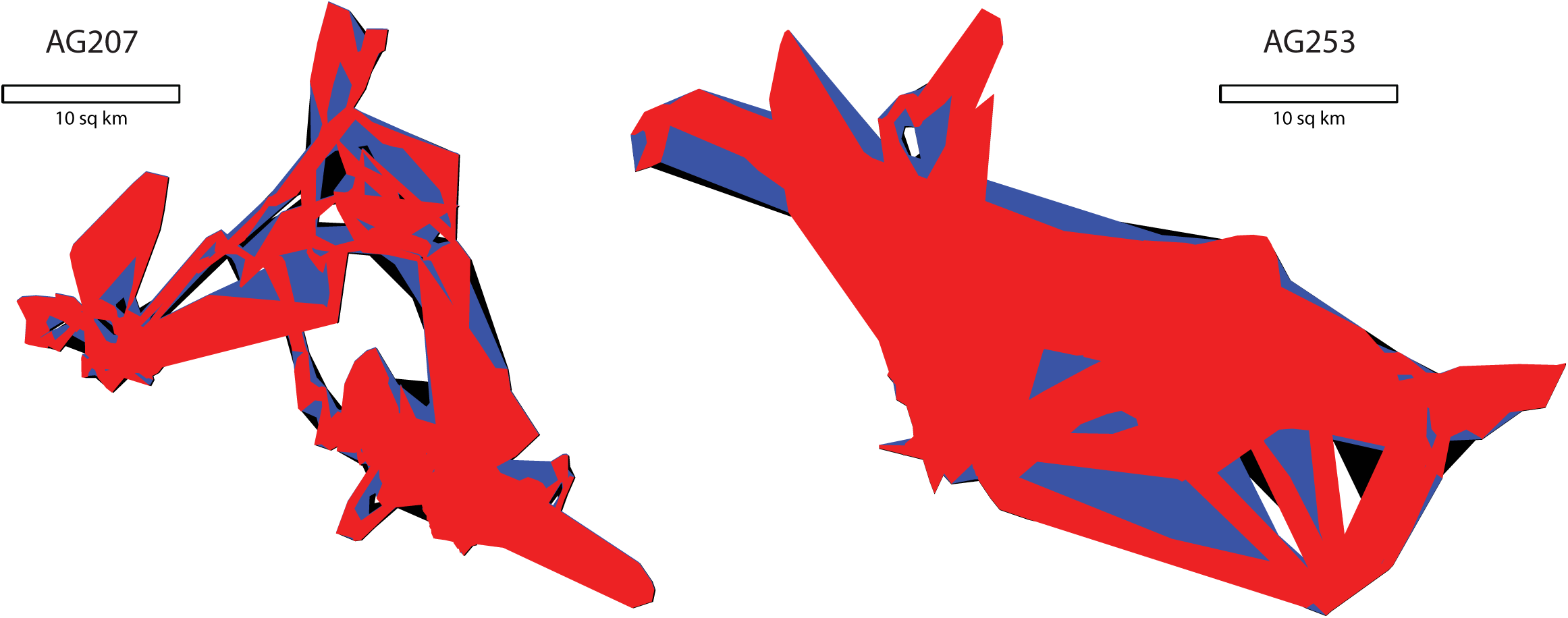
Comparison of Resulting Home Ranges. An illustration of two sets of home ranges that result from the parameter sets chosen by the algorithm (red), the low range of the guide (blue), and the high range of the guide (black). The home range set on the left is based on the sample points from the springbok AG207, and the largest home range covers 317.98 *km*^2^. The home range set on the right is based on the GPS fixes from zebra AG253, and the largest home range covers 513.06 *km*^2^.

**Figure 4.**
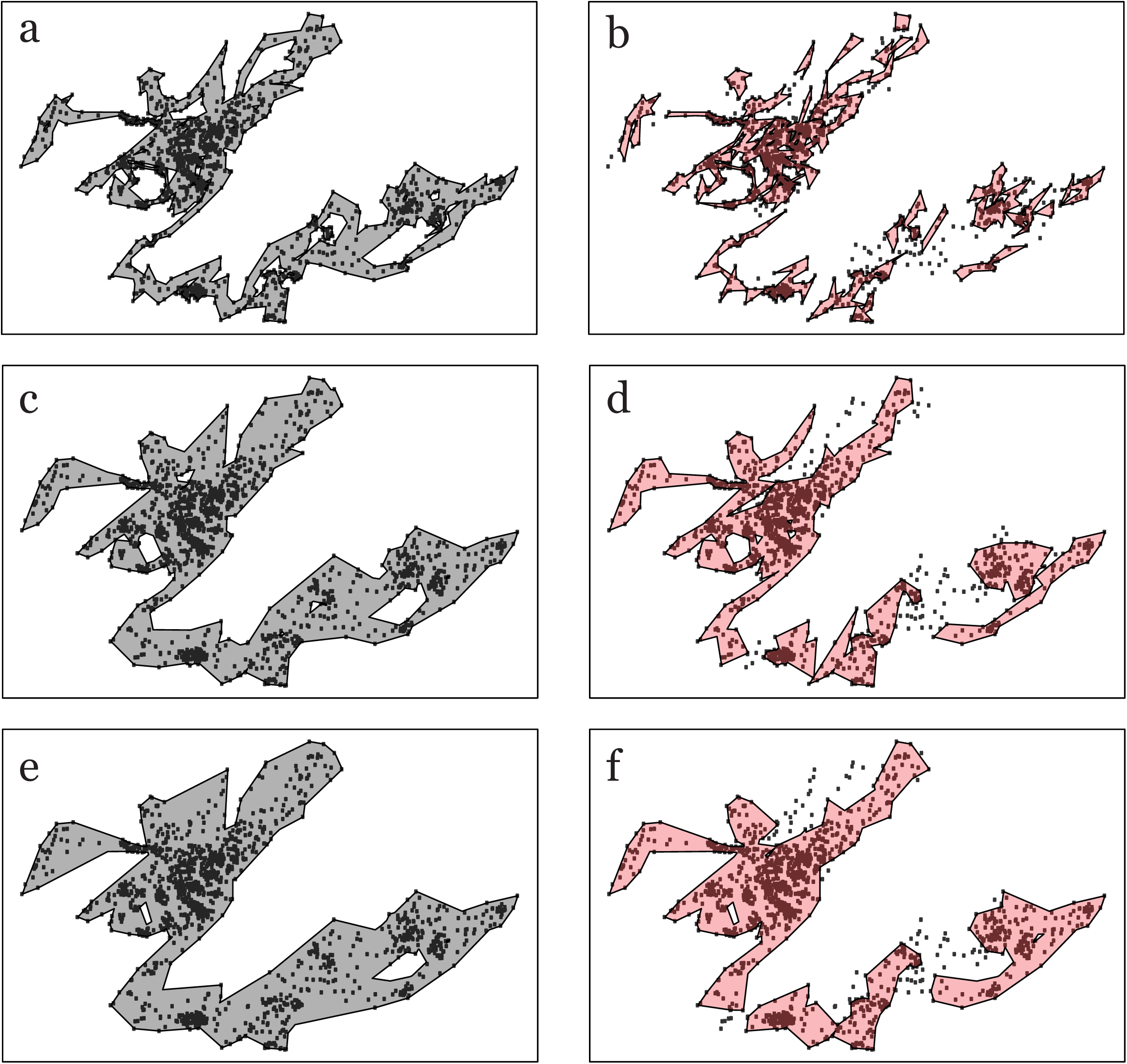
Hulls versus Isopleths. Using the simulated movement trajectory, home ranges can be delimited using the hulls themselves (a,c,e) or the isopleths (b,d,f) derived from the level of overlap among hulls (in this case, the 95% isopleth is displayed). When the *k* value is relatively small (*k*=5), the hulls (a) outline the movements of the animal very closely, offering insight, not only into core areas, but also potentially important movement corridors. Using isopleths (b) at low *k* values may result in large holes throughout the home range while failing to capture corridors. At moderate and high *k* values (c,d,e,f), both the hulls and isopleths begin to fill in many of the ancillary features, delimiting similar home ranges at slightly different rates (i.e., at *k*=25, the isopleths (f) resemble the home range outlined by the hulls at *k*=15 (c)). This illustrates the issue of underfitting when using hulls at high *k* values and overfitting when using isopleths at low *k* values. The algorithm proposed here serves to balance these two scenarios as effectively as possible.

## 5 Conclusion

Here we present a unifying protocol for parameter selection based on a cross-validation approach. Using the hulls created by the T-LoCoH method as the guiding element for choosing appropriate *s* and *k* values, one can maximize the information content of the home range, penalizing parameter sets that resemble the uninformative MCP while maintaining a level of generality that allows for inference beyond the telemetry points themselves. This approach enables consistent comparisons among the derived metrics of different individuals and species, as well as among different time periods, removing subjectivity from the T-LoCoH parameter selection process. The lack of a unifying conception of the home range contributes to the broad and inconsistent application of the term throughout the movement ecology literature and beyond. While the method proposed here has its own assumptions, it offers an objective alternative that can be applied across taxa and study sites to unify results. Ultimately, standardization will facilitate a more explicit connection between animal movement and our conception of space use patterns with major implications for the conservation and management of wildlife.

## Acknowledgements

The authors thank Perry de Valpine for his aid in developing the algorithm. We would also like to acknowledge Andy Lyons for creating the T-LoCoH package and continuing to improve it daily. Finally, we would like to thank the members of the Getz and Brashares Labs at UC Berkeley for their comments and suggestions throughout the development and writing process, particularly Dana Seidel and Briana Abrahms.

## Declarations

### Funding

The case study presented here used GPS movement data from zebra and springbok from Etosha National Park, Namibia, which were collected under a grant obtained by WMG (NIH GM083863). In addition, partial funding for this study was provided by NIH 1R01GM117617-01 to JKB and WMG. The funders had no role in study design, data collection and analysis, nor manuscript writing.

### Availability of data and materials

Please contact Wayne M. Getz for data requests.

### Author’s contributions

ERD, JKB, and WMG conceived of the project. ERD and CJC developed the algorithm. ERD ran analyses on real and simulated movement paths. ERD and CJC contributed to writing. JKB and WMG offered extensive edits to the writing and guidance throughout.

### Competing interests

The authors declare that they have no competing interests.

### Consent for publication

Not applicable.

### Ethics approval and consent to participate

All movement data were collected according to the animal handling protocol AUP R217-0509B (University of California, Berkeley).

## Additional Files

Additional file 1 — R Code for the parameter selection algorithm The code is parallelized so that multiple movement paths can be analyzed simultaneously on a multi-core computer.

